# Origin of Novel Coronavirus (COVID-19): A Computational Biology Study using Artificial Intelligence

**DOI:** 10.1101/2020.05.12.091397

**Authors:** Thanh Thi Nguyen, Mohamed Abdelrazek, Dung Tien Nguyen, Sunil Aryal, Duc Thanh Nguyen, Sandeep Reddy, Quoc Viet Hung Nguyen, Amin Khatami, Edbert B. Hsu, Samuel Yang

## Abstract

Origin of the COVID-19 virus (SARS-CoV-2) has been intensely debated in the scientific community since the first infected cases were detected in December 2019. The disease has caused a global pandemic, leading to deaths of thousands of people across the world and thus finding origin of this novel coronavirus is important in responding and controlling the pandemic. Recent research results suggest that bats or pangolins might be the hosts for SARS-CoV-2 based on comparative studies using its genomic sequences. This paper investigates the SARS-CoV-2 origin by using artificial intelligence (AI) and raw genomic sequences of the virus. More than 300 genome sequences of COVID-19 infected cases collected from different countries are explored and analysed using unsupervised clustering methods. The results obtained from various AI-enabled experiments using clustering algorithms demonstrate that all examined SARS-CoV-2 genomes belong to a cluster that also contains bat and pangolin coronavirus genomes. This provides evidence strongly supporting scientific hypotheses that bats and pangolins are probable hosts for SARS-CoV-2. At the whole genome analysis level, our findings also indicate that bats are more likely the hosts for the COVID-19 virus than pangolins.

## 1. Introduction

The COVID-19 pandemic has rapidly spread across many countries and disturbed lives of millions of people around the globe. There have been approximately 148.5 million confirmed cases of COVID-19 globally, including more than 3.1 million deaths, reported to the World Health Organization (WHO) at the end of April 2021 (WHO, 2021). Studies on understanding the virus, which was named severe acute respiratory syndrome coronavirus 2 (SARS-CoV-2), are important to propose appropriate intervention strategies and contribute to the therapeutics and vaccine development. Finding origin of SARS-CoV-2 is crucial as it helps to understand where the virus comes from via its evolutionary relationships with other biological organisms and species. This will facilitate the process of identifying and isolating the source and preventing further transmissions to the human population. This will also help to understand the outbreak dynamics, leading to the creation of informed plans for public health responses (WHO, 2020). Origin of SARS-CoV-2 however is a controversial topic with some uncertainty even after the WHO investigation commenced in October 2020 (Mallapaty, 2020).

A study by Wu et al. (2020) using a complete genome obtained from a patient who was a worker at a seafood market in Wuhan city, Hubei province, China shows that the virus is closely related to a group of SARS-like CoVs that were pre-viously found present in bats in China. It is believed that bats are the most likely reservoir hosts for SARS-CoV-2 as it is very similar to a bat coronavirus. These results are supported by a separate study by Lu et al. (2020) using genome sequences acquired from nine COVID-19 patients who were among early cases in Wuhan, China. Outcomes of a phylogenetic analysis suggest that the virus belongs to the genus *Betacoronavirus*, sub-genus *Sarbecovirus*, which includes many bat SARS-like CoVs and SARS CoVs. A study in Zhu et al. (2020) confirms this finding by analysing genomes obtained from three adult patients admitted to a hospital in Wuhan on December 27, 2019. Likewise, Zhou et al. (2020) advocate a probable bat origin of SARS-CoV-2 by using complete genome sequences of five patients at the beginning of the outbreak in Wuhan, China. One of these sequences shows 96.2% similarity to a genome sequence of a coronavirus, denoted RaTG13, which was previously obtained from a *Rhinolophus affinis* bat found in Yunnan province of China. Zhang and Holmes (2020) also highlight a similarity of about 85% between SARS-CoV-2 and RaTG13 in their receptor binding domain, which is an important region of the viral genomes for binding the viruses to the human angiotensin-converting enzyme 2 receptor.

In another study, Lam et al. (2020) found two related lineages of CoVs in pangolin genome sequences sampled in Guangxi and Guangdong provinces in China, which have similar ge-nomic organizations to SARS-CoV-2. That study suggests that pangolins could be possible hosts for SARS-CoV-2 although they are solitary animals in an endangered status with relatively small population sizes. These findings are corroborated by Zhang et al. (2020) who assembled a pangolin CoV draft genome using a reference-guided scaffolding approach based on contigs taxonomically annotated to SARS-CoV-2, SARS-CoV, and bat SARS-like CoV. Xiao et al. (2020) furthermore suggest that SARS-CoV-2 may have been formed by a recombination of a pangolin CoV-like virus with one similar to RaTG13, and that pangolins are potentially the intermediate hosts for SARS-CoV-2. On the other hand, by analysing genomic features of SARS-CoV-2, i.e. mutations in the receptor binding domain portion of the spike protein and distinct backbone of the virus, Andersen et al. (2020) determined that this novel coronavirus originated through natural processes rather than through a laboratory manipulation.

This paper applies artificial intelligence (AI)-based unsupervised clustering methods to a dataset of SARS-CoV-2 genomic sequences to provide quantitative evidence on the origin of the virus. We propose the use of hierarchical clustering algorithm and density-based spatial clustering of applications with noise (DBSCAN) method for this purpose. Two pairwise evolutionary distances between sequences based on the Jukes-Cantor method and the maximum composite likelihood method are used to enable the execution of the clustering methods. Using unsupervised clustering methods, we have been able to analyse a large dataset including 334 genomes of SARS-CoV-2 collected from different countries. Clustering results suggest that 1) SARS-CoV-2 belongs to the Sarbecovirus sub-genus of the Betacoronavirus genus, 2) bats and pangolins may have served as the hosts for SARS-CoV-2, and 3) bats are more likely the original hosts for SARS-CoV-2 than pangolins. The findings of this research provide more insights about SARS-CoV-2 and thus facilitate the progress on discovering medicines and vaccines to mitigate its impacts and prevent a similar pandemic in the future.

## 2. Materials and Methods

We downloaded 334 complete genome sequences of SARS-CoV-2 available from the GenBank database, which is maintained by the National Center for Biotechnology Information (NCBI), in early April 2020. Among these sequences, 258 were reported from USA, 49 were from China and the rest were distributed through various countries from Asia to Europe and South America. Detailed distribution of these genome sequences across 16 countries is presented in Table 1.Most of reference sequences, e.g. ones within the *Alphacoronavirus* and *Betacoronavirus* genera, are also downloaded from the NCBI GenBank and Virus-Host DB (https://www.genome.jp/virushostdb/) that covers NCBI Reference Sequences (RefSeq, release 99, March 2, 2020). Genome sequences of Guangxi pangolin CoVs (Lam et al., 2020) are downloaded from the GISAID database (https://www.gisaid.org) with accession numbers EPI_ISL_410538 - EPI_ISL_410543. A Guangdong pangolin CoV genome (Xiao et al., 2020) is also downloaded from GISAID with accession number EPI_ISL_410721. We employ three sets of reference sequences in this study with details presented in Tables 2-4. The selection of reference genomes at different taxonomic levels is based on a study in Randhawa et al. (2020) that uses the AI-based supervised decision tree method to classify novel pathogens, which include SARS-CoV-2 sequences. We aim to traverse from higher to lower taxonomic levels in searching for the SARS-CoV-2 origin by discovery of its genus, sub-genus taxonomy and its closest genome sequences.

**Table 1:** Number of COVID-19 sequences collected from different countries

| Countries | Number of sequences | Countries | Number of sequences |
| --- | --- | --- | --- |
| USA | 258 | India | 2 |
| China | 49 | Brazil | 1 |
| Japan | 5 | Italy | 1 |
| Spain | 4 | Peru | 1 |
| Taiwan | 3 | Nepal | 1 |
| Vietnam | 2 | South Korea | 1 |
| Israel | 2 | Australia | 1 |
| Pakistan | 2 | Sweden | 1 |

**Table 2:** Reference viruses from major virus classes at a high taxonomic level - Set 1

| Virus (Accession Number) | Taxonomy | Virus (Accession Number) | Taxonomy |
| --- | --- | --- | --- |
| Human adenovirus D8 (AB448767) | Adenoviridae | Murine leukemia virus (AB187566) | Ortervirales |
| TT virus sle1957 (AM711976) | Anelloviridae | Human papillomavirus type 69 (AB027020) | Papillomaviridae |
| Staphylococcus phage S13' (AB626963) | Caudovirales | Adeno-associated virus - 6 (AF028704) | Parvoviridae |
| Chili leaf curl virus-Oman (KF229718) | Geminiviridae | Cotesia plutellae polydnavirus (AY651828) | Polydnaviridae |
| Meles meles fecal virus (JN704610) | Genomoviridae | Aves polyomavirus 1 (AF118150) | Polyomaviridae |
| Chlamydia phage 3 (AJ550635) | Microviridae | Middle East respiratory syndrome (MERS) CoV (NC_019843) | Riboviria |

Unsupervised clustering methods are employed to cluster datasets comprising both query sequences (SARS-CoV-2) and reference sequences into clusters. In this paper, we propose the use of *hierarchical clustering algorithm* (Rokach and Maimon, 2005) and *density-based spatial clustering of applications with noise (DBSCAN) method* (Ester et al., 1996) for this purpose. With these two methods, we perform two steps to observe the clustering results that lead to interpretations about the taxonomy and origin of SARS-CoV-2. In the first step, we apply clustering algorithms to cluster the set of reference sequences only, and then use the same settings (i.e. values of parameters) of clustering algorithms to cluster a dataset that merges reference sequences and SARS-CoV-2 sequences. Through this step, we can find out reference sequences by which SARS-CoV-2 sequences form a group with. In the second step, we vary the settings of the clustering algorithms and observe changes in the clustering outcomes. With the second step, we are able to discover the closest reference sequences to the SARS-CoV-2 sequences and compare the similarities between genomes.

In the hierarchical clustering method, the cut-off parameter C plays as a threshold in defining clusters and thus C is allowed to change during our experiments. With regard to the DBSCAN method, the neighbourhood search radius parameter *ε* and the minimum number of neighbours parameter, which is required to identify a core point, are crucial in partitioning observations into clusters. In our experiments, we set the minimum number of neighbours to 3 and allow only the search radius parameter *ε* to vary. Outputs of the DBSCAN method may also include outliers, which are normally labelled as cluster “-1”. To facilitate the execution of the clustering methods, we propose the use of pairwise distances between sequences based on the *Jukes-Cantor method* (Jukes et al., 1969) *and the maximum composite likelihood method* (Tamura et al., 2004). The Jukes-Cantor method estimates evolutionary distances by the maximum likelihood approach using the number of substitutions between two sequences. With nucleotide sequences, the distance is defined as d = −3/4 *ln(1 − p* 4/3) where p is the ratio between the number of positions where the substitution is to a different nucleotide and the number of positions in the sequences. On the other hand, the maximum composite likelihood method considers the sum of log-likelihoods of all pairwise distances in a distance matrix as a composite likelihood because these distances are correlated owing to their phylogenetic relationships. Tamura et al. (2004) showed that estimates of pairwise distances and their related substitution parameters such as those of the Tamura-Nei model (Tamura and Nei, 1993) can be obtained accurately and efficiently by maximizing this composite likelihood. The unweighted pair group method with arithmetic mean (UPGMA) method is applied to create hierarchical cluster trees, which are used to construct dendrogram plots for the hierarchical clustering method. The UPGMA method is also employed to generate phylogenetic trees in order to show results of the DBSCAN algorithm.

## 3. Results and Discussion

In this section, we report results of hierarchical clustering algorithm and DBSCAN, each using either of the two evolutionary distance methods, i.e. the Jukes-Cantor distance and the maximum composite likelihood distance.

### 3.1. Results obtained by using the Jukes-Cantor distance

We start the experiments to search for taxonomy and origin of SARS-CoV-2 with the first set of reference genome sequences (Set 1 in Table 2). This set consists of much more diversified viruses than the other two sets (Sets 2 and 3 in Tables 3 and 4) as it includes representatives from major virus classes at the highest available virus taxonomic level. With a large coverage of various types of viruses, the use of this reference set minimizes the probability of missing out any known virus types. Outcomes of the hierarchical clustering and DB-SCAN methods are presented in Figs. 1 and 2, respectively. In these experiments, we use 16 SARS-CoV-2 sequences representing 16 countries in Table 1 for demonstration purpose. The first released SARS-CoV-2 genome of each country is selected for these experiments. Clustering outcomes on all 334 sequences are similar to those reported here. Both clustering methods consistently demonstrate that SARS-CoV-2 sequences form a cluster with a representative virus of Riboviria among 12 major virus classes (*Adenoviridae, Anelloviridae, Caudovirales, Geminiviridae, Genomoviridae, Microviridae, Ortervirales, Papillomaviridae, Parvoviridae, Polydnaviridae, Polyomaviridae*, and *Riboviria*). The Middle East respiratory syndrome (MERS) CoV, which caused the MERS outbreak in 2012, is chosen as a representative of the Riboviria realm. In hierarchical clustering (Fig. 1), when combined with reference genomes, SARS-CoV-2 genomes do not create a new cluster on their own but form a cluster with the MERS CoV, i.e. cluster “8”. With the DBSCAN method (Fig. 2), SARS-CoV-2 genomes also do not create their own cluster but form the cluster “1” with the MERS CoV. These clustering results suggest that SARS-CoV-2 belongs to the Riboviria realm.

**Table 3:** Reference viruses within the Riboviria realm - Set 2

| Virus (Accession Number) | Taxonomy | Virus (Accession Number) | Taxonomy |
| --- | --- | --- | --- |
| Grapevine rupestris stem pitting-associated virus (GRSPV) 1 (AF057136) | Betaflexiviridae | Lymantria dispar cypovirus 14 (AF389452) | Reoviridae |
| Cucumber mosaic virus (AJ276479) | Bromoviridae | Hybrid snakehead virus (KC519324) | Rhabdoviridae |
| Chiba virus (AB042808) | Caliciviridae | Rice tungro spherical virus (NC_001632) | Secoviridae |
| Mercadeo virus (NC_027819) | Flaviviridae | Bulbul CoV HKU11-796 (FJ376620) | Coronaviridae; DeltaCoV |
| Bunyamwera virus (NC_001925) | Peribunyaviridae | Avian infectious bronchitis virus (AIBV) (AY646283) | Coronaviridae; GammaCoV |
| Rice grassy stunt tenuivirus (NC_002323) | Phenuiviridae | Human CoV NL63 (NC_005831) | Coronaviridae; AlphaCoV |
| Theiler's-like virus of rats (AB090161) | Picornaviridae | SARS CoV BJ01 (AY278488) | Coronaviridae; BetaCoV |
| Turnip mosaic virus (AB194796) | Potyviridae |  |  |

**Table 4:** Reference viruses in the genus AlphaCoV and BetaCoV - Set 3

| Virus (Accession Number) | Taxonomy | Virus (Accession Number) | Taxonomy |
| --- | --- | --- | --- |
| Transmissible gastroenteritis virus (TGEV) (NC_038861) | AphaCoV | SARS CoV BJ01 (AY278488) | BetaCoV; Sarbecovirus |
| Mink CoV WD1127 (NC_023760) | AlphaCoV | Bat SARS CoV RsSHC014 (KC881005) | BetaCoV; Sarbecovirus |
| Porcine epidemic diarrhea virus (PEDV) (NC_003436) | AlphaCoV | Bat SARS CoV WIV1 (KF367457) | BetaCoV; Sarbecovirus |
| Rhinolophus bat CoV HKU2 (NC_009988) | AlphaCoV | Bat SARS CoV Rp3 (DQ071615) | BetaCoV; Sarbecovirus |
| Human CoV 229E (NC_002645) | AlphaCoV | Bat SARS CoV Rs672/2006 (FJ588686) | BetaCoV; Sarbecovirus |
| Human CoV NL63 (NC_005831) | AlphaCoV | Bat SARS CoV Rf1 (DQ412042) | BetaCoV; Sarbecovirus |
| Human CoV OC43 (NC_006213) | BetaCoV; Embecovirus | Bat SARS CoV Longquan-140 (KF294457) | BetaCoV; Sarbecovirus |
| Murine hepatitis virus (MHV) (AC_000192) | BetaCoV; Embecovirus | Bat SARS CoV HKU3-1 | BetaCoV; Sarbecovirus |
| Rousettus bat CoV HKU9 (NC_009021) | BetaCoV; Nobecovirus | Bat SARS CoV ZXC21 (MG772934) | BetaCoV; Sarbecovirus |
| Rousettus bat CoV GCCDC1 (NC_030886) | BetaCoV; Nobecovirus | Bat SARS CoV ZC45 (MG772933) | BetaCoV; Sarbecovirus |
| MERS CoV (NC_019843) | BetaCoV; Merbecovirus | Bat CoV RaTG13 (MN996532) | BetaCoV; Sarbecovirus |
| Pipistrellus bat CoV HKU5 (NC_009020) | BetaCoV; Merbecovirus | Guangxi pangolin CoV GX/P4L (EPI_ISL_410538) | BetaCoV; Sarbecovirus |
| Tylonycteris bat CoV HKU4 (NC_009019) | BetaCoV; Merbecovirus | Guangxi pangolin CoV GX/P1E (EPI_ISL_410539) | BetaCoV; Sarbecovirus |
| Bat Hp-betaCoV/Zhejiang2013 (NC_025217) | BetaCoV; Hibecovirus | Guangxi pangolin CoV GX/P5L (EPI_ISL_410540) | BetaCoV; Sarbecovirus |
| SARS CoV BtKY72 (KY352407) | BetaCoV; Sarbecovirus | Guangxi pangolin CoV GX/P5E (EPI_ISL_410541) | BetaCoV; Sarbecovirus |
| Bat CoV BM48-31/BGR/2008 (GU190215) | BetaCoV; Sarbecovirus | Guangxi pangolin CoV GX/P2V (EPI_ISL_410542) | BetaCoV; Sarbecovirus |
| SARS CoV LC5 (AY395002) | BetaCoV; Sarbecovirus | Guangxi pangolin CoV GX/P3B (EPI_ISL_410543) | BetaCoV; Sarbecovirus |
| SARS CoV SZ3 (AY304486) | BetaCoV; Sarbecovirus | Guangdong pangolin CoV (EPI_ISL_410721) | BetaCoV; Sarbecovirus |
| SARS CoV Tor2 (AY274119) | BetaCoV; Sarbecovirus |  |  |

**Figure 1:**
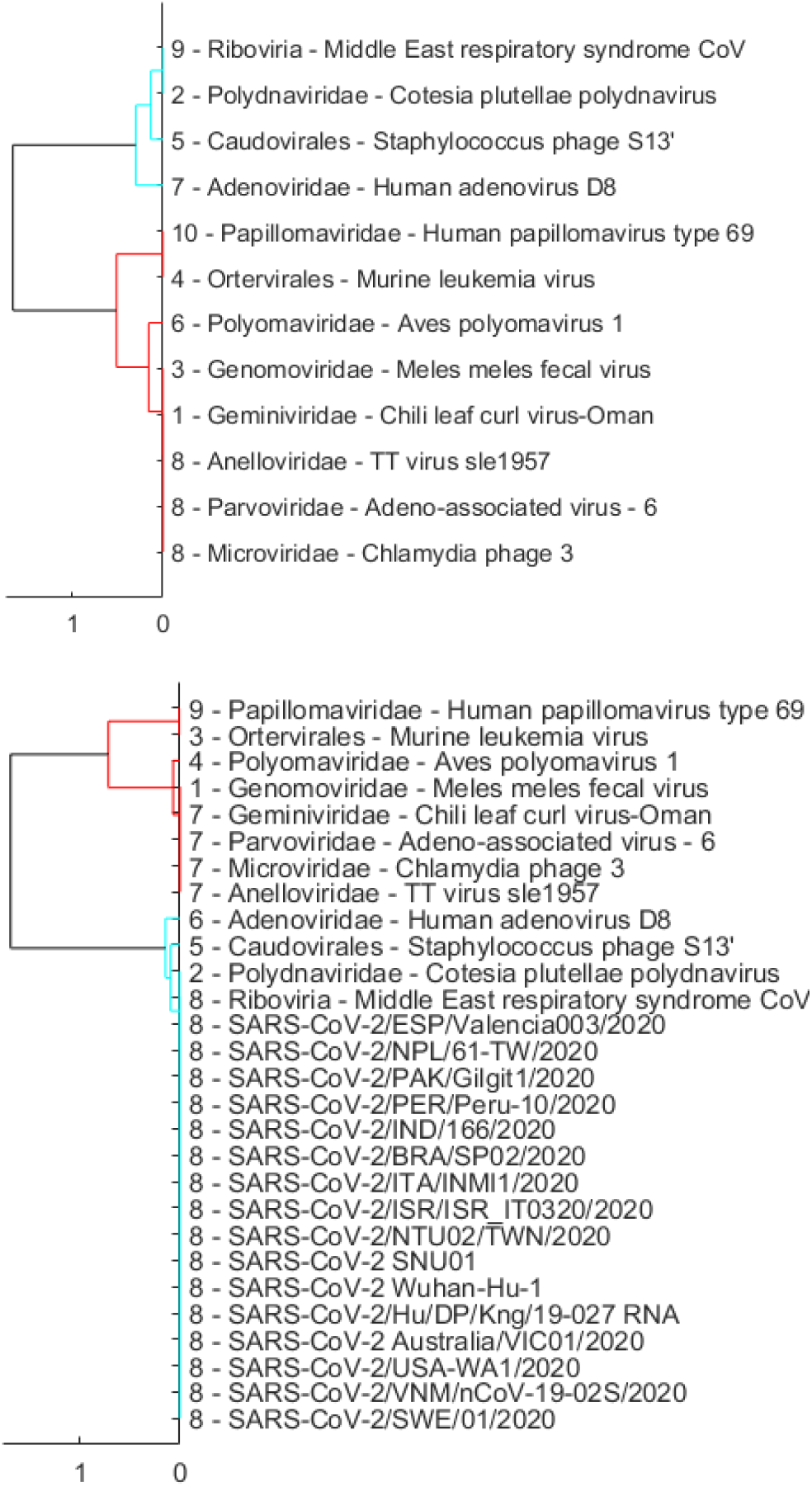
Dendrogram plots showing hierarchical clustering results using only the reference sequences in Set 1 (Table 2) with the cut-off parameter *C* equal to 5 * 10^−4^ (top), and using a set that merges 16 representative SARS-CoV-2 sequences and reference sequences with *C* also set to 5 * 10^−4^ (bottom). A number at the beginning of each virus name indicates the cluster that virus belongs to after clustering.

**Figure 2:**
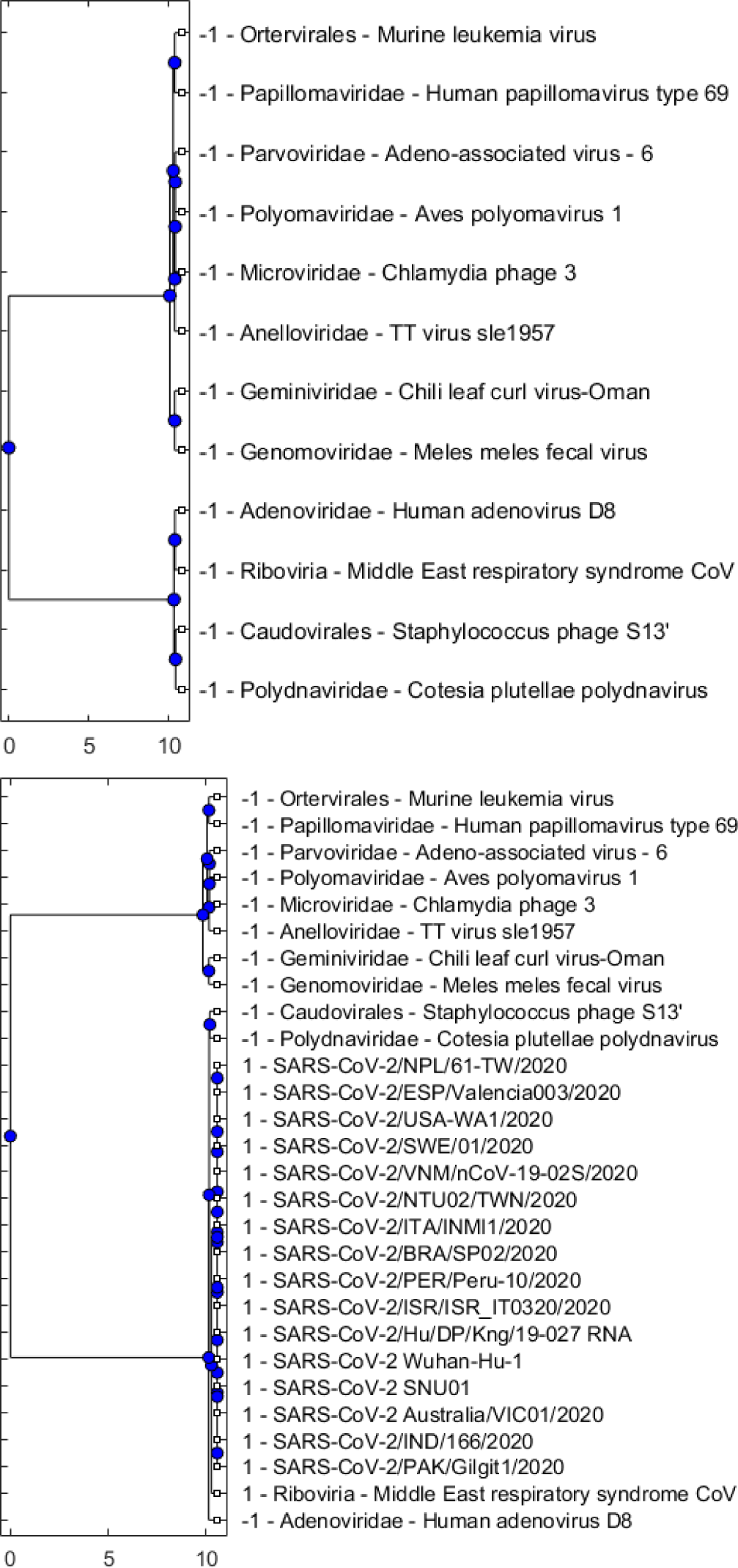
Phylogenetic trees showing DBSCAN results using only the reference sequences in Set 1 (Table 2) with the search radius parameter *ε* equal to0.7 (left), and using a set that merges SARS-CoV-2 sequences and reference sequences with *ε* also set to 0.7 (right). As Set 1 includes representatives of major virus classes and the minimum number of neighbours is set to 3 while *ε* is set to 0.7, DBSCAN considers individual viruses as outliers (top). When the dataset is expanded to include SARS-CoV-2 sequences, DBSCAN forms cluster “1” that includes all SARS-CoV-2 sequences and the MERS CoV, which represents the Riboviria realm (bottom).

Once we have been able to identify SARS-CoV-2 as belonging to the Riboviria realm, we move to the next lower taxonomic level that consists of 12 virus families within Riboviria. These families are presented in Set 2 (Table 3) that includes *Betaflexiviridae, Bromoviridae, Caliciviridae, Coronaviridae, Flaviviridae, Peribunyaviridae, Phenuiviridae, Picornaviridae, Potyviridae, Reoviridae, Rhabdoviridae*, and *Secoviridae*. Results of the two clustering methods presented in Figs. 3 and 4 show that SARS-CoV-2 sequences form a group with viruses in the *Coronaviridae* family. As we also include in Set 2 representatives of four genera in the *Coronaviridae* family (i.e. *Alpha-coronavirus - AlphaCoV, Betacoronavirus - BetaCoV, Delta-coronavirus, Gammacoronavirus*), we are able to look further to the next taxonomic level within this experiment and observe that SARS-CoV-2 belongs to the *Betacoronavirus* genus (see Figs. 3 and 4).

**Figure 3:**
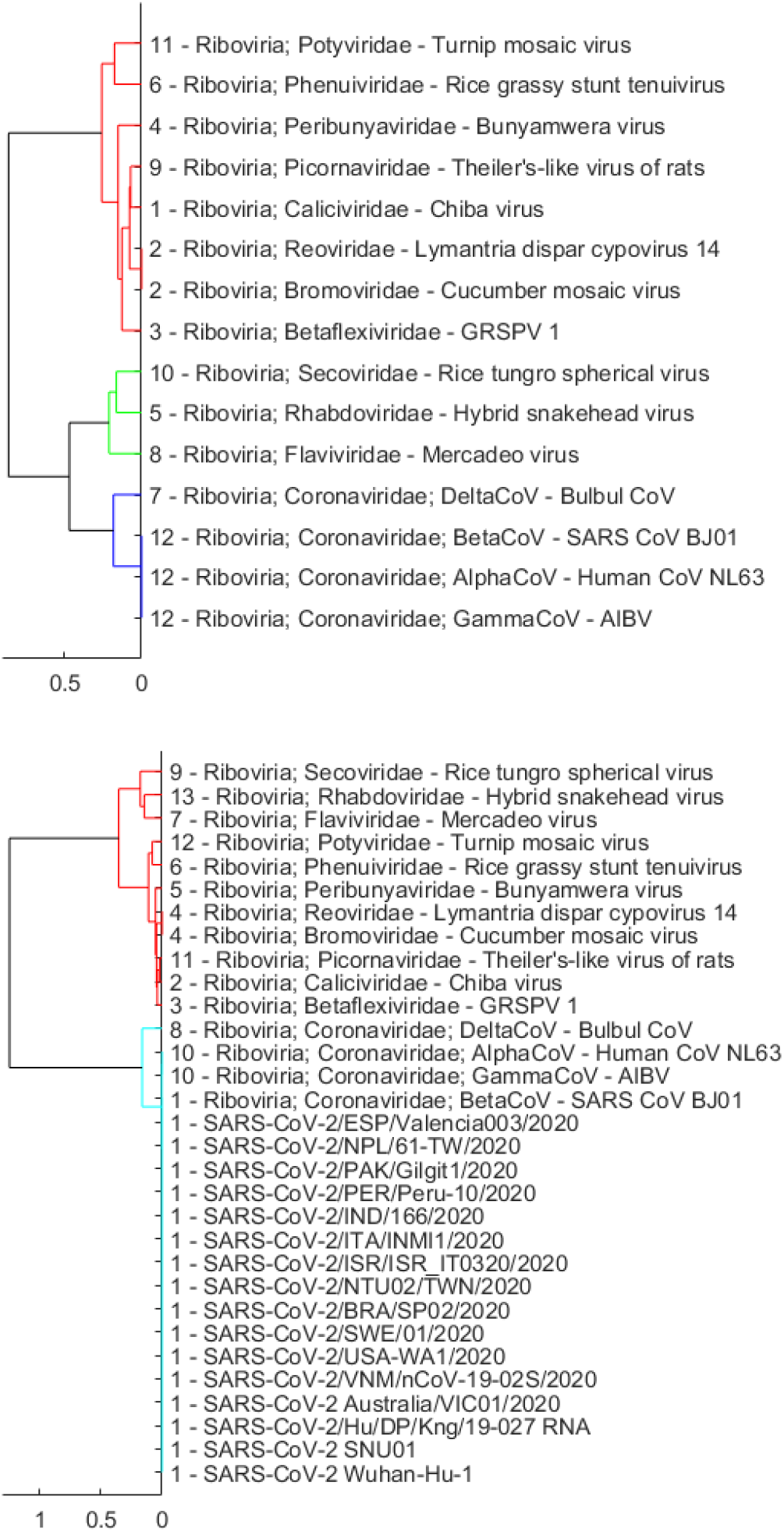
Dendrogram plots showing hierarchical clustering results using only the reference sequences in Set 2 (Table 3) with the cut-off parameter *C* equal to 0.001 (top), and using a set that merges SARS-CoV-2 sequences and reference sequences with *C* also set to 0.001 (bottom).

**Figure 4:**
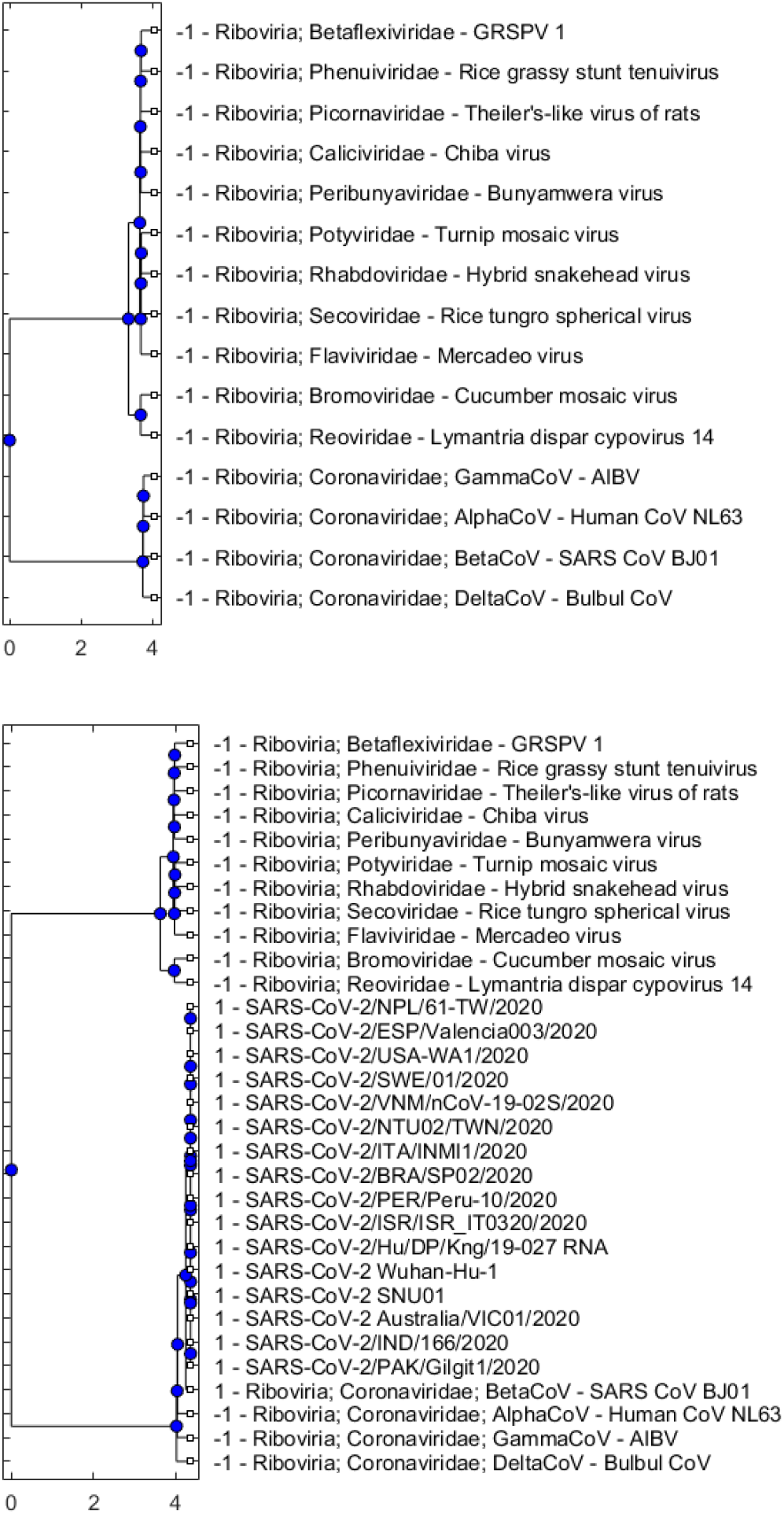
Phylogenetic trees showing DBSCAN results using only the reference sequences in Set 2 (Table 3) with the search radius parameter *ε* equal to 0.6 (top), and using a set that merges SARS-CoV-2 sequences and reference sequences with *ε* also set to 0.6 (bottom).

There are 5 sub-genera in the *Betacoronavirus* genus, including *Embecovirus, Nobecovirus, Merbecovirus, Hibecovirus, Sarbecovirus*. In the next lower taxonomic level, we investigate which of these sub-genera that SARS-CoV-2 belongs to or whether SARS-CoV-2 genomes create a new cluster on their own. We use Set 3 (Table 4) that includes 37 reference sequences for this investigation. Six representatives of the *Alpha-coronavirus* genus, which is proximal to the *Betacoronavirus* genus, are also included in Set 3. The rest of Set 3 comprises 2 viruses of the *Embecovirus* sub-genus, 2 viruses of the *Nobecovirus* sub-genus, 3 viruses of the *Merbecovirus* sub-genus, 1 virus of the *Hibecovirus* sub-genus and 23 viruses of the *Sarbecovirus* sub-genus. Most of the representatives of the *Sarbecovirus* sub-genus are SARS CoVs and bat SARS-like CoVs. Notably, we also include in this set 6 sequences of Guangxi pangolin CoVs deposited to the GISAID database by Lam et al. (2020) and a sequence of Guangdong pangolin CoV by Xiao et al. (2020).

Evolutionary distances between each of the reference genomes in Set 3 (Table 4) to the 334 SARS-CoV-2 genomes based on the Jukes-Cantor method are presented in Fig. 5. We can observe that these distances are almost constant across 334 SARS-CoV-2 sequences, which are collected in 16 countries (Table 1) over approximately 3 months since late December 2019. This implies that not much variation of SARS-CoV-2 genomes has occurred over time and across countries. As in Fig. 5, there are several groups of reference genomes shown via the closeness of the distance lines. For example, the top group contains genomes of *AlphaCoV* viruses (refer to the taxonomy in Table 4) that are quite evolutionarily divergent from SARS-CoV-2 sequences. The middle group of lines comprises most of the *BetaCoV* viruses, especially those in the *Sarbecovirus* sub-genus. The bottom lines identify reference viruses that are closest to SARS-CoV-2, which include bat CoV RaTG13, Guangdong pangolin CoV, bat SARS CoV ZC45 and bat SARS CoV ZXC21. The bat CoV RaTG13 line at the bottom is notably distinguished from other lines while the Guangdong pangolin CoV line is the second closest to SARS-CoV-2. The similarities between bat CoV RaTG13, Guang-dong pangolin CoV and Guangxi pangolin CoV GX/P4L with SARS-CoV-2/Australia/VIC01/2020, produced by the SimPlot software (Lole et al., 1999), are displayed in Fig. 6. Consistent with the results presented in Fig. 5, bat CoV RaTG13 is shown closer to SARS-CoV-2 than pangolin CoVs.

**Figure 5:**
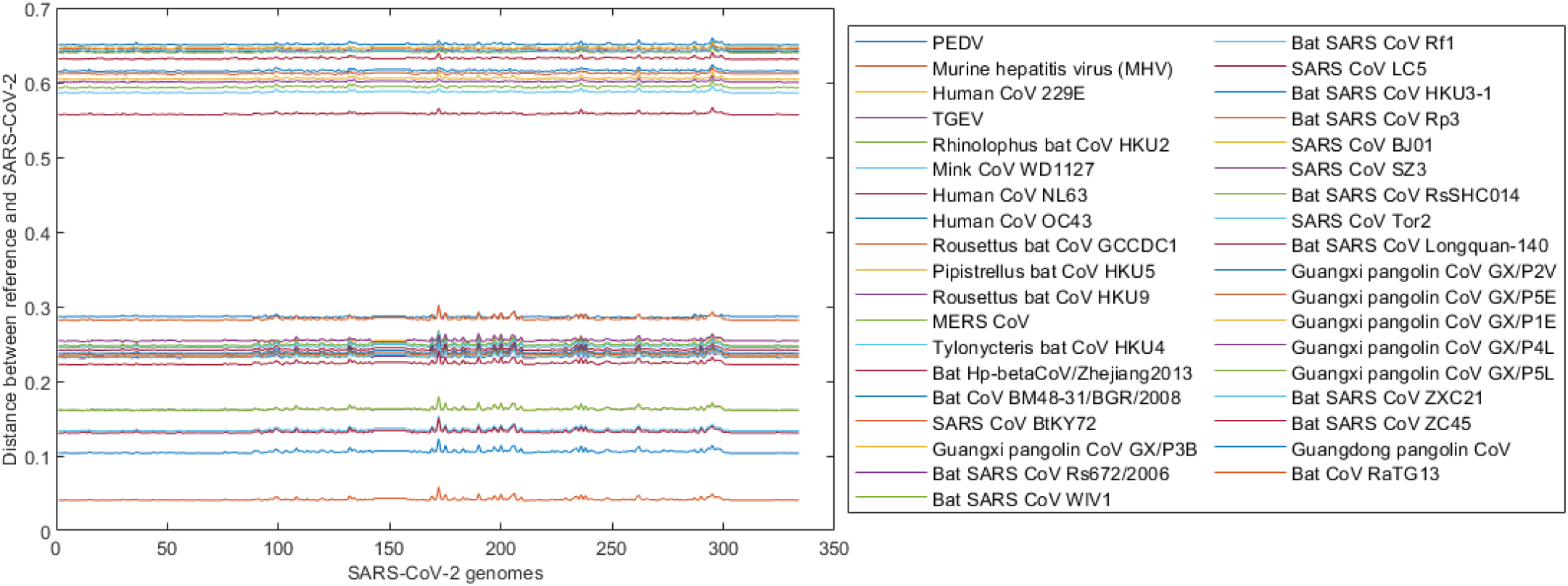
Distances between each of the reference genomes in Set 3 (Table 4) with 334 SARS-CoV-2 genomes where the latter are ordered by the released date (earliest to latest) over a period of approximately 3 months, from late December 2019 to late March 2020. These pairwise distances are computed based on the Jukes-Cantor method using whole genomes. The line at the bottom, for instance, represents the distances between the bat CoV RaTG13 genome with each of the 334 SARS-CoV-2 genomes.

**Figure 6:**
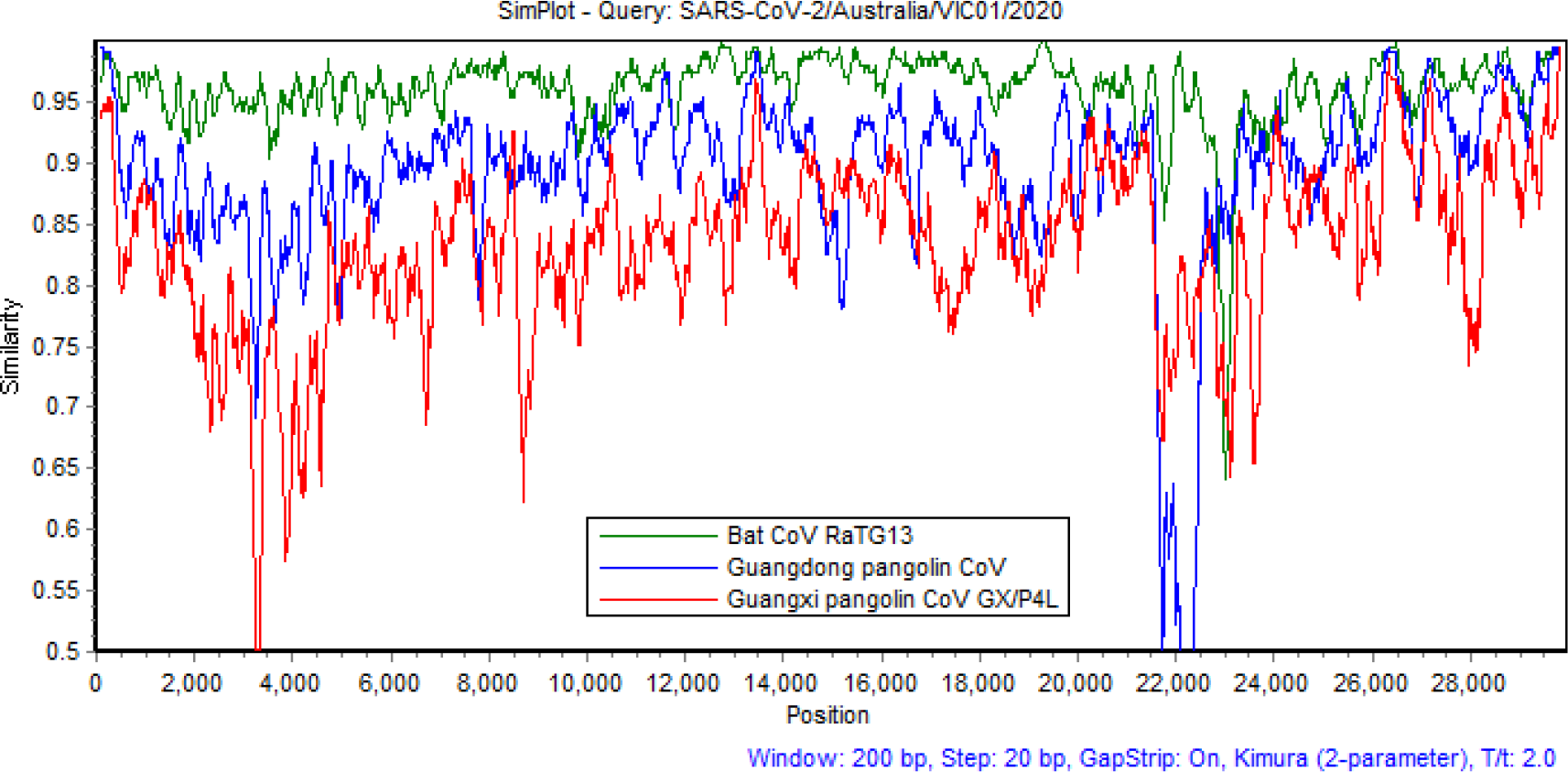
Similarities between genomes of bat CoV RaTG13, Guangdong pangolin CoV and Guangxi pangolin CoV GX/P4L with the SARS-CoV-2/Australia/VIC01/2020 genome sequence.

Fig. 7 shows outcomes of the hierarchical clustering method using Set 3 of reference sequences in Table 4. When the cut-off parameter *C* is set equal to 0.7, the hierarchical clustering algorithm separates the reference sequences into 6 clusters in which cluster “5” comprises all examined viruses of the *Sarbecovirus* sub-genus, including many SARS CoVs, bat SARS-like CoVs and pangolin CoVs (Fig. 7A). It is observed that the algorithm reasonably groups viruses into clusters, for example, the genus *AlphaCoV* is represented by cluster “4” while the sub-genera *Embecovirus, Nobecovirus, Merbecovirus*, and *Hibecovirus* are labelled as clusters “3”, “6”, “2”, and “1”, respectively. Using the same cut-off value of 0.7, we next perform clustering on a dataset that merges reference sequences and 16 representative SARS-CoV-2 sequences (see Fig. 7B). Results on all 334 SARS-CoV-2 sequences are similar to those on the 16 representative sequences. The outcome presented in Fig. 7B shows that the 16 representative SARS-CoV-2 sequences fall into cluster “5”, which comprises the *Sarbecovirus* sub-genus. The number of clusters is still 6 and the membership structure of the clusters is the same as in the case of clustering reference sequences only

**Figure 7:**
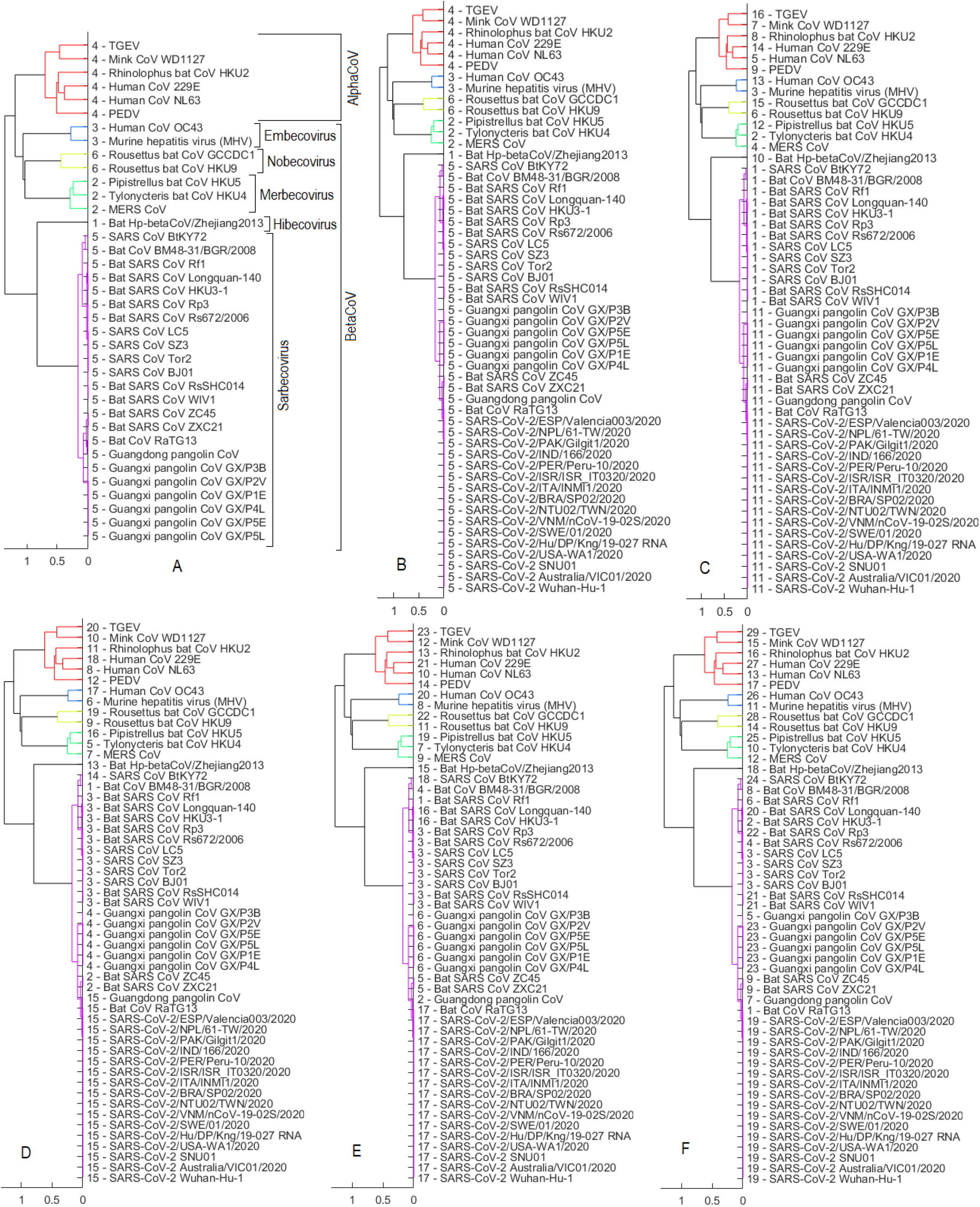
Set 3 of reference sequences - results obtained by using hierarchical clustering via dendrogram plots. Numbers at the beginning of each virus label indicate the cluster that virus is a member of as a result of the clustering algorithm. (A) when the cut-off *C* is equal to 0.7 and using Set 3 of reference sequences only: there are 6 clusters where cluster “5” covers all examined viruses in the *Sarbecovirus* sub-genus. (B) when cut-off *C* is still equal to 0.7 and the dataset now merges between reference sequences and 16 representative SARS-CoV-2 sequences (merged set). (C) using the merged set with *C* = 0.1. (D) using the merged set with *C* = 0.03. (E) using the merged set with *C* = 0.01. (F) using the merged set with *C* = 0.001.

(Fig. 7A), except that the *Sarbecovirus* cluster now has been expanded to also contain SARS-CoV-2 sequences. By comparing Figs. 7A and 7B, we believe that SARS-CoV-2 is naturally part of the *Sarbecovirus* sub-genus. This realization is substantiated by moving to Fig. 7C that shows a clustering outcome when the cut-off parameter *C* is decreased to 0.1. In Fig. 7C, while 3 members of the *Merbecovirus* sub-genus (i.e. *Pipistrellus* bat CoV HKU5, *Tylonycteris* bat CoV HKU4 and MERS CoV) are divided into 3 clusters (”12”, “2” and “4”) or members of the *Sarbecovirus* cluster separate themselves into 2 clusters “’1” and “11”, sequences of SARS-CoV-2 still join the cluster “11’ with other members of *Sarbecovirus* such as 3 bat viruses (bat SARS CoV ZC45, bat SARS CoV ZXC21, bat CoV RaTG13) and 7 pangolin CoVs.

As the cut-off parameter *C* decreases, the number of clusters increases. This is an expected outcome because the cut-off threshold line moves closer to the leaves of the dendrogram. When the cut-off *C* is reduced to 0.03 (Fig. 7D), there are only 2 viruses (bat CoV RaTG13 and Guangdong pangolin CoV) that can form a cluster with SARS-CoV-2 (labelled as cluster “15”). These are 2 viruses closest to SARS-CoV-2 based on the whole genome analysis. Results in Figs. 7C and 7D therefore provide evidence that bats or pangolins could be possible hosts for SARS-CoV-2. We next reduce the cut-off *C* to 0.01 as in Fig. 7E. At this stage, only bat CoV RaTG13 is within the same cluster with SARS-CoV-2 (cluster “17”). We thus believe that bats are the more probable hosts for SARS-CoV-2 than pangolins. The inference of our AI-enabled analysis is in line with a result in Cárdenas-Conejo et al. (2020) that investigates the polyprotein 1ab of SARS-CoV-2 and suggests that this novel coronavirus has more likely been arisen from viruses infecting bats rather than pangolins. When the cut-off *C* is reduced to 0.001 as in Fig. 7F, we observe that the total number of clusters now increases to 29 and more importantly, SARS-CoV-2 sequences do not combine with any other reference viruses but form its own cluster “19”. Could we use this clustering result (Fig. 7F) to infer that SARS-CoV-2 might not originate in bats or pangolins? This is a debatable question because the answer depends on the level of details we use to differentiate between the species or organisms. The cut-off parameter in hierarchical clustering can be considered as the level of details. With the results obtained in Fig. 7D (and also in the experiments with the DBSCAN method presented next), we support a hypothesis that bats or pangolins are the probable origin of SARS-CoV-2. This is because we observe considerable similarity between SARS-CoV-2 and bat CoV RaTG13 (or Guangdong pangolin CoV) compared to the similarity between viruses that originated in the same host. For example, bat SARS-like CoVs such as bat SARS CoV Rf1, bat SARS CoV Longquan-140, bat SARS CoV HKU3-1, bat SARS CoV Rp3, bat SARS CoV Rs672/2006, bat SARS CoV RsSHC014, bat SARS CoV WIV1, bat SARS CoV ZC45 and bat SARS CoV ZXC21 had the same bat origin. In Fig. 7D, these viruses however are separated into 2 different clusters (”3” and “2”) while all 16 SARS-CoV-2 representatives are grouped together with bat CoV RaTG13 and Guangdong pangolin CoV in cluster “15”. This demonstrates that the difference between the same origin viruses (e.g. bat SARS CoV WIV1 and bat SARS CoV ZC45) is larger than the difference between SARS-CoV-2 and bat CoV RaTG13 (or Guangdong pangolin CoV). Therefore, SARS-CoV-2 is deemed to have very likely originated in the same host with bat CoV RaTG13 or Guangdong pangolin CoV, which is bat or pangolin, respectively.

Clustering outcomes of the DBSCAN method via phylogenetic trees using Set 3 of reference sequences (Table 4) are presented in Fig. 8. We first apply DBSCAN to reference sequences only, which results in 3 clusters and several outliers (Fig. 8A). The search radius parameter *ε* is set equal to 0.55. As we set the minimum number of neighbours parameter to 3, it is expected that viruses of the sub-genera *Embecovirus, Nobecovirus* and *Hibecovirus* are detected as outliers “-1” because there are only 1 or 2 viruses in these sub-genera. Three viruses of the *Merbecovirus* sub-genus (i.e. *Tylonycteris* bat CoV HKU4, *Pipistrellus* bat CoV HKU5 and MERS CoV) are grouped into the cluster “2”. All examined viruses of the *Sarbecovirus* sub-genus are joined in cluster “1” while the *Alpha-CoV* viruses are combined into cluster “3”. Fig. 8B shows an outcome of DBSCAN with the same *ε* value of 0.55 and the dataset has been expanded to include 16 representative SARS-CoV-2 sequences. We observe that genomes of SARS-CoV-2 fall into the cluster “1”, which includes all the examined *Sarbecovirus* viruses. When *ε* is decreased to 0.3 in Fig. 8C, all members of the *Merbecovirus* cluster or the *AlphaCoV* cluster become outliers while 16 SARS-CoV-2 genomes still stick with the *Sarbecovirus* cluster. In line with the results obtained by using hierarchical clustering in Fig. 7, those obtained in Fig. 8B and 8C using the DBSCAN method give us the confidence to confirm that SARS-CoV-2 is part of the *Sarbecovirus* subgenus. Fig. 8D shows that bat CoV RaTG13 and Guangdong pangolin CoV are closest to SARS-CoV-2 as they join with 16 SARS-CoV-2 representatives in cluster “2”. This again substantiates the probable bat or pangolin origin of SARS-CoV-2. By reducing *ε* to 0.1 as in Fig. 8E, the Guangdong pangolin CoV becomes an outlier whilst SARS-CoV-2 sequences form a cluster (”3”) with only bat CoV RaTG13. This further confirms our findings when using the hierarchical clustering in Fig. 7 that bats are more likely the hosts for the SARS-CoV-2 than pangolins. When *ε* is decreased to 0.01 as in Fig. 8F, SARS-CoV-2 genomes form its own cluster “3”, which is separated with any bat or pangolin genomes. As with the result in Fig. 7F by the hierarchical clustering, this result also raises a question whether SARS-CoV-2 really originated in bats or pangolins. In Fig. 8D, it is again observed that the similarity between SARS-CoV-2 and bat CoV RaTG13 (or Guangdong pangolin CoV) is larger than the similarity between bat SARS CoVs, which share the same bat origin. Specifically, SARS-CoV-2, bat CoV RaTG13 and Guangdong pangolin CoV are grouped together in cluster “2” while bat SARS CoVs are divided into 2 clusters, i.e. bat SARS CoV ZXC21 and bat SARS CoV ZC45 are in cluster “2” whereas other bat SARS CoVs are in cluster “3”. We thus suggest that SARS-CoV-2 probably has the same origin with bat CoV RaTG13 or Guangdong pangolin CoV. In other words, bats or pangolins are the probable origin of SARS-CoV-2.

**Figure 8:**
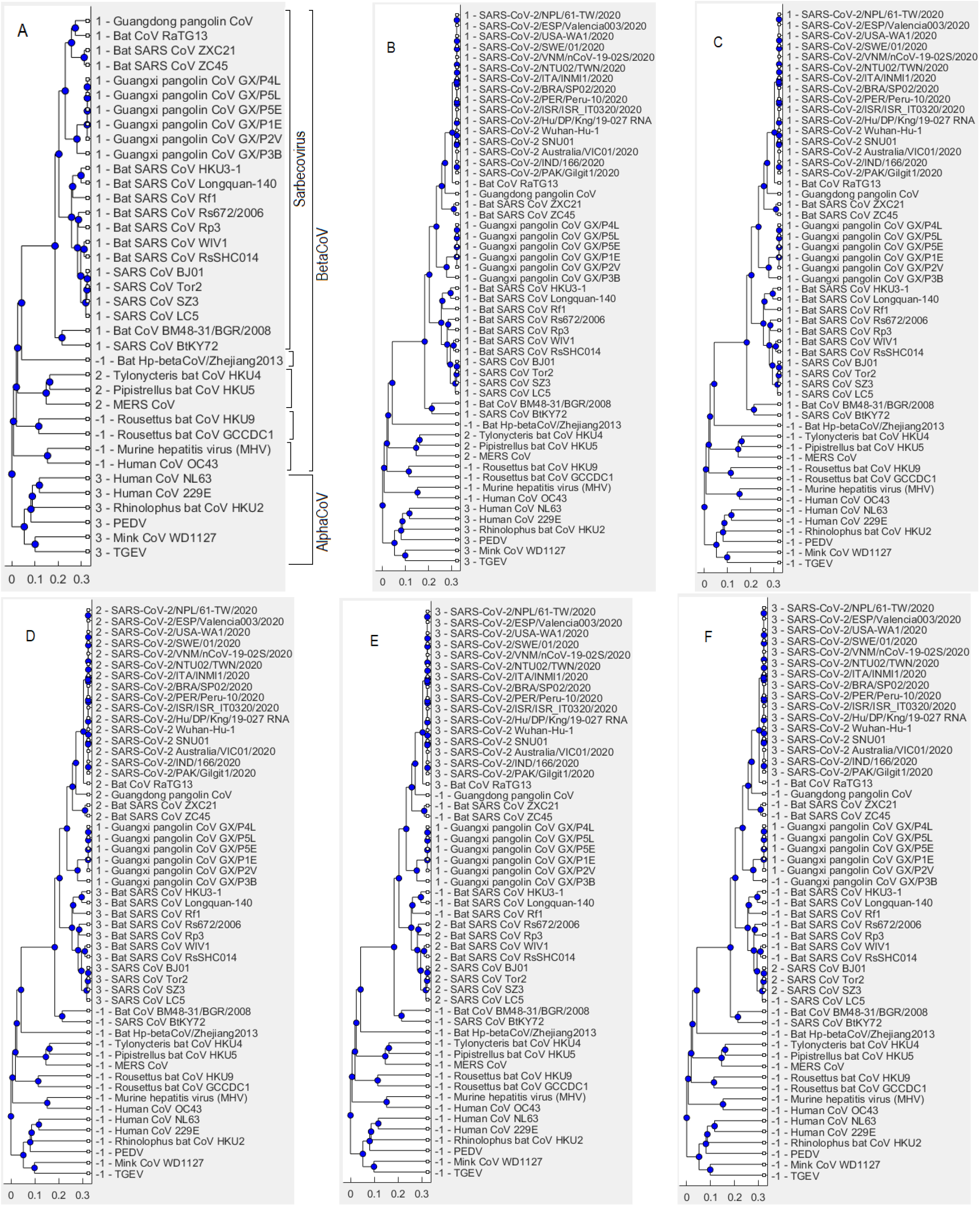
Set 3 of reference sequences - results obtained by using DBSCAN via phylogenetic trees. Numbers at the beginning of each virus label indicate the cluster that virus is a member of as a result of the clustering algorithm. (A) when the search radius parameter *ε* is equal to 0.55 and using Set 3 of reference sequences only: there are 3 clusters where the cluster “1” covers all examined viruses in the *Sarbecovirus* sub-genus. (B) when search radius *ε* is still equal to 0.55 and the dataset now merges between reference sequences and 16 representative SARS-CoV-2 sequences (merged set). (C) using the merged set with *ε* = 0.3. (D) using the merged set with *ε* = 0.15. (E) using the merged set with *ε* = 0.1. (F) using the merged set with *ε* = 0.01.

All results presented above are obtained using the pairwise distances estimated by the *Jukes-Cantor method*. The following subsection reports clustering results obtained using evolutionary distances calculated by the *maximum composite likelihood method*.

### 3.2. Results obtained by using the maximum composite likelihood distance method

This subsection presents results of two clustering methods, i.e. hierarchical clustering and DBSCAN, using the sequence distances computed by the maximum composite likelihood method (Tamura et al., 2004), which was conducted in the MEGA X software (Kumar et al., 2018). These results are greatly similar to those obtained by using the Jukes-Cantor distance method shown throughout the paper. In these experiments, the clustering methods are applied to a dataset that combines reference sequences in Set 3 (Table 4) and 16 representative genomes of 16 countries in Table 1. When a country has more than one collected genome, the first released genome of that country is selected for this experiment. Fig. 9 demonstrates the distances estimated by the maximum composite likelihood method between each of the reference sequences and 16 representative SARS-CoV-2 genomes. The lines are almost parallel indicating that SARS-CoV-2 genome is not altered much across countries, which is in line with the results obtained using the Jukes-Cantor distance estimates in Fig. 5. The bat CoV RaTG13 is again shown much closer to SARS-CoV-2 than pangolin CoVs and other reference viruses although the distance range in Fig. 9 is larger than that in Fig. 5, i.e. [0, ~1.6] versus [0, 0. ∼7], respectively.

**Figure 9:**
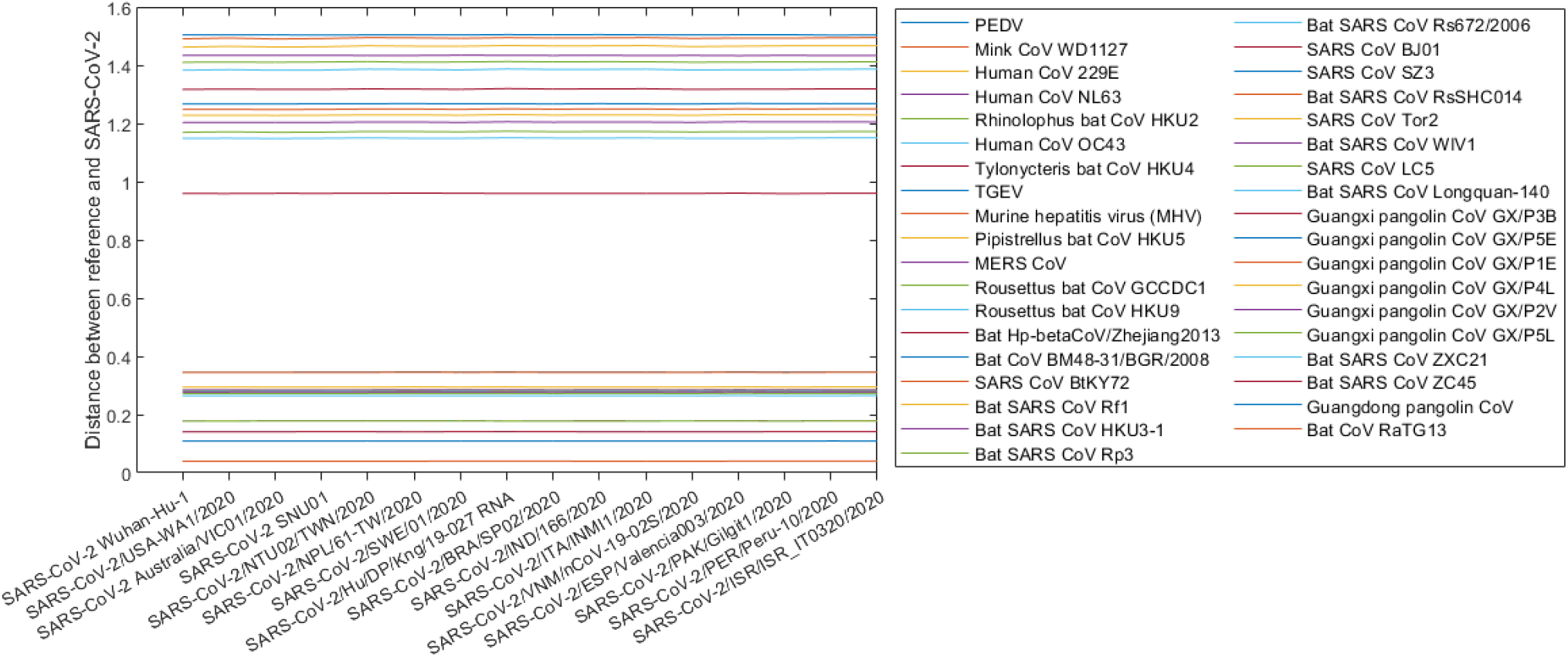
Evolutionary distances between reference genomes and 16 representative SARS-CoV-2 genomes of 16 countries based on the maximum composite likelihood method. Each line is composed of 16 data points representing 16 pairwise distances. The bottom line, for instance, shows the distances between bat CoV RaTG13 and each of 16 representative genomes on the *x*-axis.

In Fig. 10A, when the hierarchical clustering cut-off parameter is set equal to 0.1, all 16 representative SARS-CoV-2 genomes are grouped into cluster “12”, which also includes other viruses of the *Sarbecovirus* sub-genus of the *BetaCoV* genus. When moving from Fig. 10A to Fig. 10B, even though members of the *Sarbecovirus* cluster (”12” in Fig. 10A) are split into 2 clusters “1” and “2” in Fig. 10B, the SARS-CoV-2 sequences are still grouped into cluster “14” with other members of the *Sarbecovirus* sub-genus such as bat CoV RaTG13, Guangdong pangolin CoV, bat SARS CoV ZXC21 and bat SARS CoV ZC45. These results provide us with a confidence on confirming the *Sarbecovirus* sub-genus of the SARS-CoV-2. This is consistent with the result based on the Jukes-Cantor distances shown in Fig. 7. Fig. 10C shows that SARS-CoV-2 genomes are combined only with that of bat CoV RaTG13 when the cut-off parameter is decreased to 0.001. This again indicates that bats are the more likely origin of SARS-CoV-2 than pangolins. When we reduce the cut-off parameter to 0.0001, the SARS-CoV-2 sequences create their own cluster “22” and this questions the probable bat or pangolin origin of SARS-CoV-2. However, in Fig. 10B, we also find that the similarity between SARS-CoV-2 and bat CoV RaTG13 (or Guangdong pangolin CoV) is larger than the similarity between viruses having the same origin. For example, bat SARS CoV WIV1 and bat SARS CoV ZC45 have the same bat origin but they are divided into 2 clusters (”1” and “14”) while all 16 SARS-CoV-2 representatives are grouped into cluster “14” with bat CoV RaTG13 and Guangdong pangolin CoV. This implies that SARS-CoV-2 may have originated in bats or pangolins.

**Figure 10:**
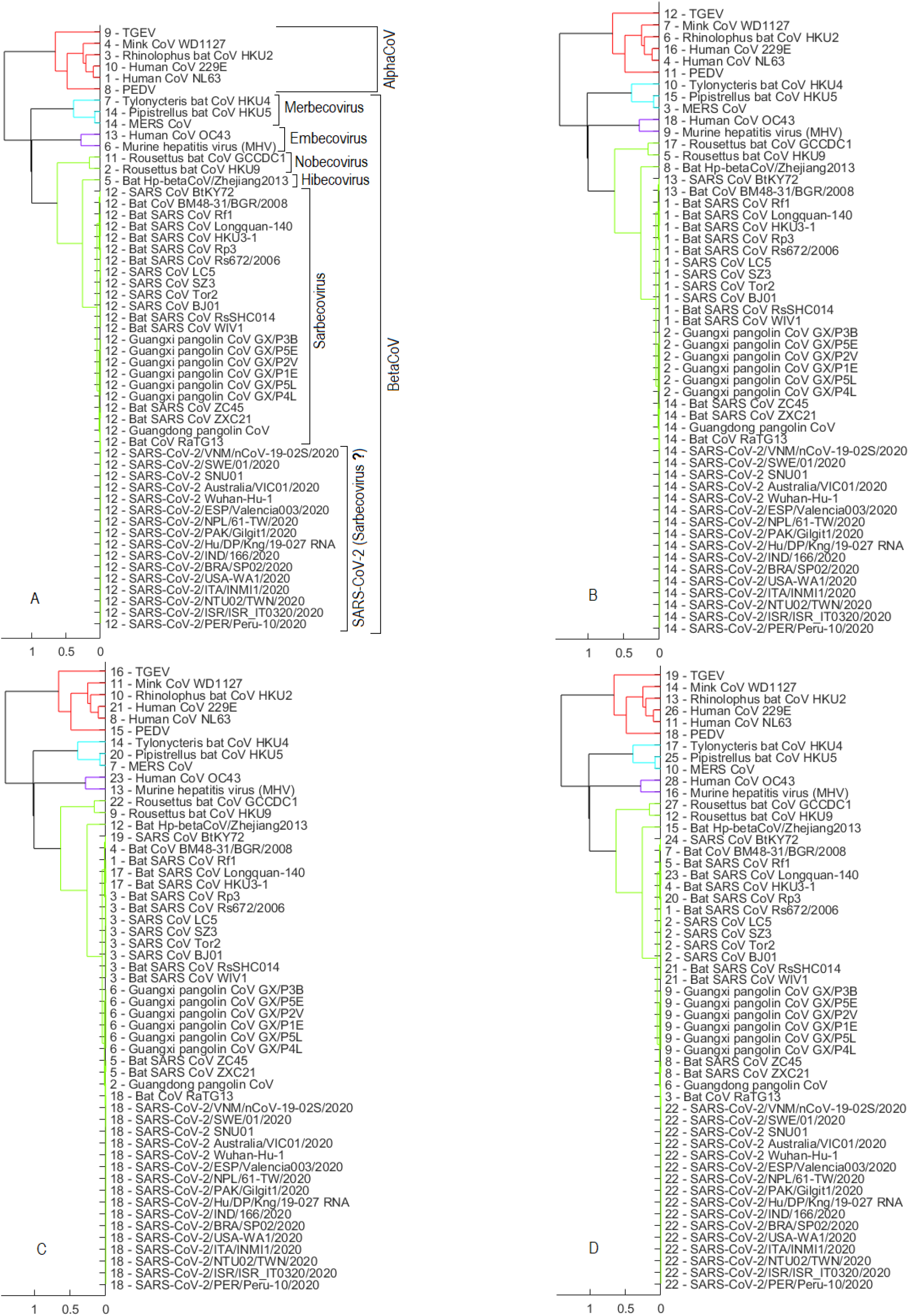
Hierarchical clustering results using pairwise distances based on the maximum composite likelihood method, (A) the cut-off parameter is set to 0.1, (B) cut-off equal to 0.01, (C) cut-off is 0.001, and (D) cut-off is 0.0001.

Results of the DBSCAN algorithm using distances estimated by the maximum composite likelihood method are similar to those obtained by the Jukes-Cantor distance method in Fig. 8. In summary, these AI-based quantitative results using the unsupervised hierarchical clustering and DBSCAN methods provide consistent evidence to suggest that 1) SARS-CoV-2 belongs to the *Sarbecovirus* sub-genus of the *Betacoronavirus* genus, 2) bats and pangolins may have served as the hosts for SARS-CoV-2, and 3) bats are the more probable origin of SARS-CoV-2 than pangolins.

## 4. Conclusions

The severity of COVID-19 pandemic has initiated a race in finding origin of SARS-CoV-2. Studies on genome sequences obtained from early patients in Wuhan city in China suggest the probable bat origin of the virus based on similarities between these sequences and those obtained from bat CoVs previously reported in China. Other studies afterwards found that SARS-CoV-2 genome sequences are also similar to pangolin CoV sequences and accordingly raised a hypothesis on the pangolin origin of SARS-CoV-2. This paper has investigated origin of SARS-CoV-2 using *two unsupervised clustering algorithms with two evolutionary distance estimation methods*, and more than 300 raw genome sequences of SARS-CoV-2 collected from various countries around the world. Outcomes of these AI-enabled methods are analysed, leading to a confirmation on the *Coronaviridae* family of SARS-CoV-2. More specifically, the SARS-CoV-2 belongs to the sub-genus *Sarbecovirus* within the genus *Betacoronavirus* that includes SARS-CoV, which caused the global SARS pandemic in 2002-2003 (Drosten et al., 2003; Wolfe et al., 2007). The results of various clustering experiments show that SARS-CoV-2 genomes are more likely to form a cluster with the bat CoV RaTG13 genome than pangolin CoV genomes, which were constructed from samples collected in Guangxi and Guangdong provinces in China. This indicates that bats are more likely the hosts for SARS-CoV-2 than pangolins.

The findings of this research on the large dataset of 334 SARS-CoV-2 genomic sequences provide more insights about SARS-CoV-2 and thus facilitate the progress on discovering medicines and vaccines to mitigate its impacts and prevent a similar pandemic in future. This study among many AI studies in the challenging battle against the COVID-19 pandemic (Nguyen et al., 2020) has shown the power and capabilities of AI, especially from the computational biology and medicine perspective. The race to produce effective treatment drugs and vaccines is ongoing and much needed in the fight against the COVID-19 pandemic. Further study in this direction is strongly encouraged by a recent success of AI in identifying powerful new kinds of antibiotic from a pool of more than 100 million molecules (Stokes et al., 2020). With the capability of analyzing large datasets and extracting knowledge in an intelligent and efficient manner, discovery of newer therapeutics and vaccine strategies using AI is becoming ever more realistic (Etzioni, 2020).

## References

1. Andersen, K.G., Rambaut, A., Lipkin, W.I., Holmes, E.C., Garry, R.F., 2020. The proximal origin of sars-cov-2. Nature Medicine 26, 450–452.

2. Cárdenas-Conejo, Y., Liñan-Rico, A., García-Rodríguez, D.A., Centeno-Leija, S., Serrano-Posada, H., 2020. An exclusive 42 amino acid signature in pp1ab protein provides insights into the evolutive history of the 2019 novel human-pathogenic coronavirus (sars-cov-2). Journal of Medical Virology 92, 688–692.

3. Drosten, C., Günther, S., Preiser, W., Van Der Werf, S., Brodt, H.R., Becker, S., Rabenau, H., Panning, M., Kolesnikova, L., Fouchier, R.A., et al., 2003. Identification of a novel coronavirus in patients with severe acute respiratory syndrome. New England Journal of Medicine 348, 1967–1976.

4. Ester, M., Kriegel, H.P., Sander, J., Xu, X., et al., 1996. A density-based algorithm for discovering clusters in large spatial databases with noise., in: KDD, pp. 226–231.

5. Etzioni, O., 2020. AI can help scientists find a covid-19 vaccine. URL: https://www.wired.com/story/opinion-ai-can-help-find-scientists-find-a-covid-19-vaccine/.

6. Jukes, T.H., Cantor, C.R., et al., 1969. Evolution of protein molecules. Mammalian Protein Metabolism 3, 21–132.

7. Kumar, S., Stecher, G., Li, M., Knyaz, C., Tamura, K., 2018. Mega x: molecular evolutionary genetics analysis across computing platforms. Molecular Biology and Evolution 35, 1547–1549.

8. Lam, T.T.Y., Jia, N., Zhang, Y.W., Shum, M.H.H., Jiang, J.F., Zhu, H.C., Tong, Y.G., Shi, Y.X., Ni, X.B., Liao, Y.S., et al., 2020. Identifying sars-cov-2- related coronaviruses in malayan pangolins. Nature 583, 282–285.

9. Lole, K.S., Bollinger, R.C., Paranjape, R.S., Gadkari, D., Kulkarni, S.S., Novak, N.G., Ingersoll, R., Sheppard, H.W., Ray, S.C., 1999. Full-length human immunodeficiency virus type 1 genomes from subtype c-infected seroconverters in india, with evidence of intersubtype recombination. Journal of Virology 73, 152–160.

10. Lu, R., Zhao, X., Li, J., Niu, P., Yang, B., Wu, H., Wang, W., Song, H., Huang, B., Zhu, N., et al., 2020. Genomic characterisation and epidemiology of 2019 novel coronavirus: implications for virus origins and receptor binding. The Lancet 395, 565–574.

11. Mallapaty, S., 2020. Where did covid come from? who investigation begins but faces challenges. Nature 587, 341–342.

12. Nguyen, T.T., Nguyen, Q.V.H., Nguyen, D.T., Hsu, E.B., Yang, S., Eklund, P., 2020. Artificial intelligence in the battle against coronavirus (covid-19): a survey and future research directions. arXiv preprint 2008.07343.

13. Randhawa, G.S., Soltysiak, M.P., El Roz, H., de Souza, C.P., Hill, K.A., Kari, L., 2020. Machine learning using intrinsic genomic signatures for rapid classification of novel pathogens: Covid-19 case study. PLoS One 15, e0232391.

14. Rokach, L., Maimon, O., 2005. Clustering methods, in: Data Mining and Knowledge Discovery Handbook. Springer, pp. 321–352.

15. Stokes, J.M., Yang, K., Swanson, K., Jin, W., Cubillos-Ruiz, A., Donghia, N.M., MacNair, C.R., French, S., Carfrae, L.A., Bloom-Ackermann, Z., et al., 2020. A deep learning approach to antibiotic discovery. Cell 180, 688–702.

16. Tamura, K., Nei, M., 1993. Estimation of the number of nucleotide substitutions in the control region of mitochondrial dna in humans and chimpanzees. Molecular Biology and Evolution 10, 512–526.

17. Tamura, K., Nei, M., Kumar, S., 2004. Prospects for inferring very large phylogenies by using the neighbor-joining method. Proceedings of the National Academy of Sciences 101, 11030–11035.

18. WHO, 2020. Origin of sars-cov-2. URL: https://www.who.int/publications/i/item/origin-of-sars-cov-2.

19. WHO, 2021. Who coronavirus disease (covid-19) dashboard. URL: https://covid19.who.int/.

20. Wolfe, N.D., Dunavan, C.P., Diamond, J., 2007. Origins of major human infectious diseases. Nature 447, 279–283.

21. Wu, F., Zhao, S., Yu, B., Chen, Y.M., Wang, W., Song, Z.G., Hu, Y., Tao, Z.W., Tian, J.H., Pei, Y.Y., et al., 2020. A new coronavirus associated with human respiratory disease in china. Nature 579, 265–269.

22. Xiao, K., Zhai, J., Feng, Y., Zhou, N., Zhang, X., Zou, J.J., Li, N., Guo, Y., Li, X., Shen, X., et al., 2020. Isolation of sars-cov-2-related coronavirus from malayan pangolins. Nature 583, 286–289.

23. Zhang, T., Wu, Q., Zhang, Z., 2020. Probable pangolin origin of sars-cov-2 associated with the covid-19 outbreak. Current Biology 30, 1346–1351.

24. Zhang, Y.Z., Holmes, E.C., 2020. A genomic perspective on the origin and emergence of sars-cov-2. Cell 181, 223–227.

25. Zhou, P., Yang, X.L., Wang, X.G., Hu, B., Zhang, L., Zhang, W., Si, H.R., Zhu, Y., Li, B., Huang, C.L., et al., 2020. A pneumonia outbreak associated with a new coronavirus of probable bat origin. Nature 579, 270–273.

26. Zhu, N., Zhang, D., Wang, W., Li, X., Yang, B., Song, J., Zhao, X., Huang, B., Shi, W., Lu, R., et al., 2020. A novel coronavirus from patients with pneumonia in china, 2019. New England Journal of Medicine.

